# Temporal neural mechanisms underlying conscious access to different levels of facial stimulus contents

**DOI:** 10.1101/220426

**Authors:** Shen-Mou Hsu, Yu-Fang Yang

**Author notes:** Corresponding author:* Shen-Mou Hsu, Imaging Center for Integrated Body, Mind and Culture Research, National Taiwan University, No.49, Fanglan Rd., Da’an Dist., Taipei, 10617, Taiwan (R.O.C).

## Abstract

An important issue facing the empirical study of consciousness concerns how the contents of incoming stimuli gain access to conscious processing. According to classic theories, facial stimuli are processed in a hierarchical manner. However, it remains unclear how the brain determines which level of stimulus contents is consciously accessible when facing an incoming facial stimulus. Accordingly, with a magnetoencephalography technique, this study aims to investigate the temporal dynamics of the neural mechanism mediating which level of stimulus content is consciously accessible. Participants were instructed to view masked target faces at threshold, so that according to behavioral responses, their perceptual awareness alternated from consciously accessing facial identity in some trials to being able to consciously access facial configuration features but not facial identity in other trials. Conscious access at these two levels of facial contents were associated with a series of differential neural events. Before target presentation, different patterns of phase angle adjustment were observed between the two types of conscious access. This effect was followed by stronger phase clustering for awareness of facial identity immediately during stimulus presentation. After target onset, conscious access to facial identity, as opposed to facial configural features, was able to elicit more robust late positivity. In conclusion, we suggest that the stages of neural events, ranging from prestimulus to stimulus-related activities, may operate in combination to determine which level of stimulus contents is consciously accessed. Conscious access may thus be better construed as comprising various forms that depend on the level of stimulus contents accessed.

## NEW & NOTEWORTHY

The present study investigates that when facing an incoming facial stimulus, how does the brain determine which level of stimulus contents is consciously accessible? Using magnetoencephalography, we show that prestimulus activities together with stimulus-related activities may operate in combination to determine conscious face detection or identification. This finding is distinct from the previous notion that conscious face detection precedes identification and provides novel insights into the temporal dynamics of different levels of conscious face perception.

## INTRODUCTION

Consciousness is a hallmark feature of the human mind, yet it remains mysterious. One important issue with empirical studies of consciousness concerns the neural mechanism underlying how external information gains access to conscious processing and in turn, defines reportable conscious content (Baars 2002). Although the nature of conscious access is subject to intense investigation, previous research on this issue has commonly focused on comparing how human brains differentially respond to aware and unaware stimuli (Aru et al. 2012; Dehaene and Changeux 2011). In most of these studies, participants report their general awareness of incoming stimuli, such as whether a stimulus is seen or not, or just specify their awareness of a certain aspect of stimulus contents, such as whether a stimulus is a face or a house. However, stimuli are processed in a hierarchical manner, from very simple features to complex conglomerates, and thereby may consist of different levels of contents. For example, a face stimulus consists of at least two levels of contents from basic-level facial configural features to deep-level facial identity, and the processing of information involved in each level is distinct (Bruce and Young 1986; Haxby et al. 2000). On the one hand, facial configural features concern the arrangement of facial features (e.g., two eyes above a nose) and thereby specify the stimulus as a face. On the other hand, facial identity involves encoding subtle variations in the shape or spacing of the features and thereby specify facial differences among individuals (Maurer et al. 2002).

From this perspective, there might exist dissociable forms of conscious perception that associate with access to various levels of stimulus contents (Campana and Tallon-Baudry 2013; Hochstein and Ahissar 2002; Kouider et al. 2010; Windey et al. 2013). In the domain of temporal processing of conscious face perception, research with different experimental approaches (Navajas et al. 2013; Rodriguez et al. 2012; Sandberg et al. 2013) has consistently shown that conscious detection of the presence of faces elicits early neural responses which peak around 170 m from stimulus onset, an event-related component corresponding to the well-established N170 (Bentin et al. 1996) or M170 component (Liu et al. 2002) in the literature. In a separate line of research (Genetti et al. 2009; Tanskanen et al. 2007), neural responses associated with awareness of facial identity are found to emerge relatively late around 230 ms and this effect is followed by an increase of the P300 component. The notion that conscious face detection precedes conscious face identification is further supported by a recent study on object perception (Koivisto et al. 2017). In this study, stimulus hierarchy is taken into account by comparing electrophysiological correlates between two separate tasks, in which participants were required to detect the presence of a digit stimulus at threshold or to identify whether the digit was smaller or larger than 5. The results also indicate an early neural signature, 200-300 ms after stimulus onset, responsible for conscious detection, whereas conscious identification is correlated with a late positive enhancement of the P300 component around 400 ms. However, opposing results have also been reported, in which conscious face detection may also evoke the P300 component (Rodriguez et al. 2012) and conscious face identification is also associated with the M170 component (Tanskanen et al. 2007). More importantly, in all these studies, separate experiments and different task requirements were carried out to individually investigate conscious access to a specific level of stimulus contents. One crucial issue remains unsolved; when facing an incoming stimulus, how does the brain determine which level of stimulus contents is consciously accessible? In other words, how different levels of stimulus contents from the very same facial stimulus compete for conscious access. Furthermore, previous studies have shown that the prestimulus alpha phase may modulate subsequent aware/unaware states of visual perception (Busch et al. 2009; Mathewson et al. 2009), revealing that distinct patterns of rhythmic phase adjustment that precede the upcoming stimuli may partly determine whether a stimulus is consciously perceived. Accordingly, is it possible that conscious access to different levels of stimulus contents could also be partly resolved even before stimulus onset?

To address these issues, the present study used the hierarchical contents of facial stimuli, we briefly presented a masked face target at threshold. Based on their reports, participants were able to consciously access its facial identify in some trials (i.e., conscious face identification). In other trials, they were able to consciously access facial configuration features without recognizing its facial identity (i.e., conscious face detection). In the remaining trials, participants were unable to consciously detect the presence of the face. In this regard, participants expressed their conscious experience of different levels of stimulus contents based on the very same stimulus, but the level that was consciously accessed alternated over trials within a single paradigm. Meanwhile, magnetoencephalography (MEG) signals were recorded to reveal the temporal dynamics of conscious access. The objective of this study is to unveil the temporal neural mechanism mediating conscious access to different levels of stimulus contents when facing an incoming stimulus. Specifically, we investigated the respective roles of prestimulus and stimulus-related phase and event-related activities in the mechanism.

## METHODS

### Participants

Thirteen right-handed participants without a past neurological or psychiatric history were enrolled as participants (8 males, mean age ± STD = 26.38 ± 3.23 years, range = 21 – 31). All participants had normal or corrected-to-normal vision and provided written informed consent. This number of participants was determined based on previous MEG or electroencephalography (EEG) research on face perception (Liu et al. 2002) and consciousness (Busch et al., 2009) using similar experimental approaches. All procedures were carried out in accordance with the Declaration of Helsinki and were approved by the ethical committee of National Taiwan University.

### Stimuli

The grayscale stimuli consisted of 56 different facial images of the same target celebrity (Andy Liu) and 24 facial images of different non-target celebrities. All images were collected from the internet and depict various viewing conditions and backgrounds to reduce the possibility that the participants could use a small set of low-level features to perform the task. Most of the target images were in a frontal view and with smiles. To ensure that a few images with different gaze direction and neutral expression did not confound the results, we repeated our analysis by excluding those images and the same pattern of results could still be obtained. Additionally, 34 texture patterns were generated by scrambling randomly selected facial images into 1 x 1 pixel squares. Masks were created in a similar manner by scrambling the faces into 8 x 8 pixel squares. All the stimuli were adjusted and matched on low-level physical attributes (luminance, contrast and spatial frequency) using the SHINE toolbox (Willenbockel et al. 2010). The face images subtended a horizontal visual angle of 2.4° and a vertical angle of 2.7° around the center of the screen. Notably, only one target identity was used throughout the experiment. This strategy aimed to avoid introducing additional noise during the analysis, given that our pilot study showed that different target identities would require differential mask intensities to obtain the luminance that yielded stimuli at threshold (see below).

Prior to the study, participants provided familiarity ratings for the target celebrity without viewing the experimental stimuli. The participants indicated their familiarity with the celebrity on a scale of 1 to 3: 1) “I have not heard of this celebrity at all,” 2) “I have heard of this celebrity, but I am not familiar with the face,” or 3) “I have heard of this celebrity, and I am familiar with the face.” All participants reported a score of 3.

### Procedure

As shown in Fig. 1, each trial began with the presentation of a fixation cross for 800-1000 ms, followed by a forward mask for 300 ms, a stimulus for 17 ms, and a backward mask for 33 ms. The stimulus was either a target face (Andy Liu), the face of another celebrity, or a texture. The technique of sandwich masking, a commonly used approach for attenuating face perception, and the mask durations were all based on similar parameters employed in prior research (Kouider and Dehaene 2007). After a blank of 250 ms, a response window with three options was displayed. Participants had up to 3000 ms to report (1) that they could recognize that the stimulus was the target face by selecting the option “Liu” (written in traditional Chinese); (2) that they could not recognize whether the face was the target or could recognize that the stimulus was the non-target face by selecting the option “Face”; or (3) that they could not see a face by selecting the option “No”. The participants responded by pressing buttons with their right index, middle and ring fingers. The positions of the three response options were randomized across trials. Pressing a button initiated a new trial after the 1400-1800 ms inter-trial interval.

**Fig. 1.**
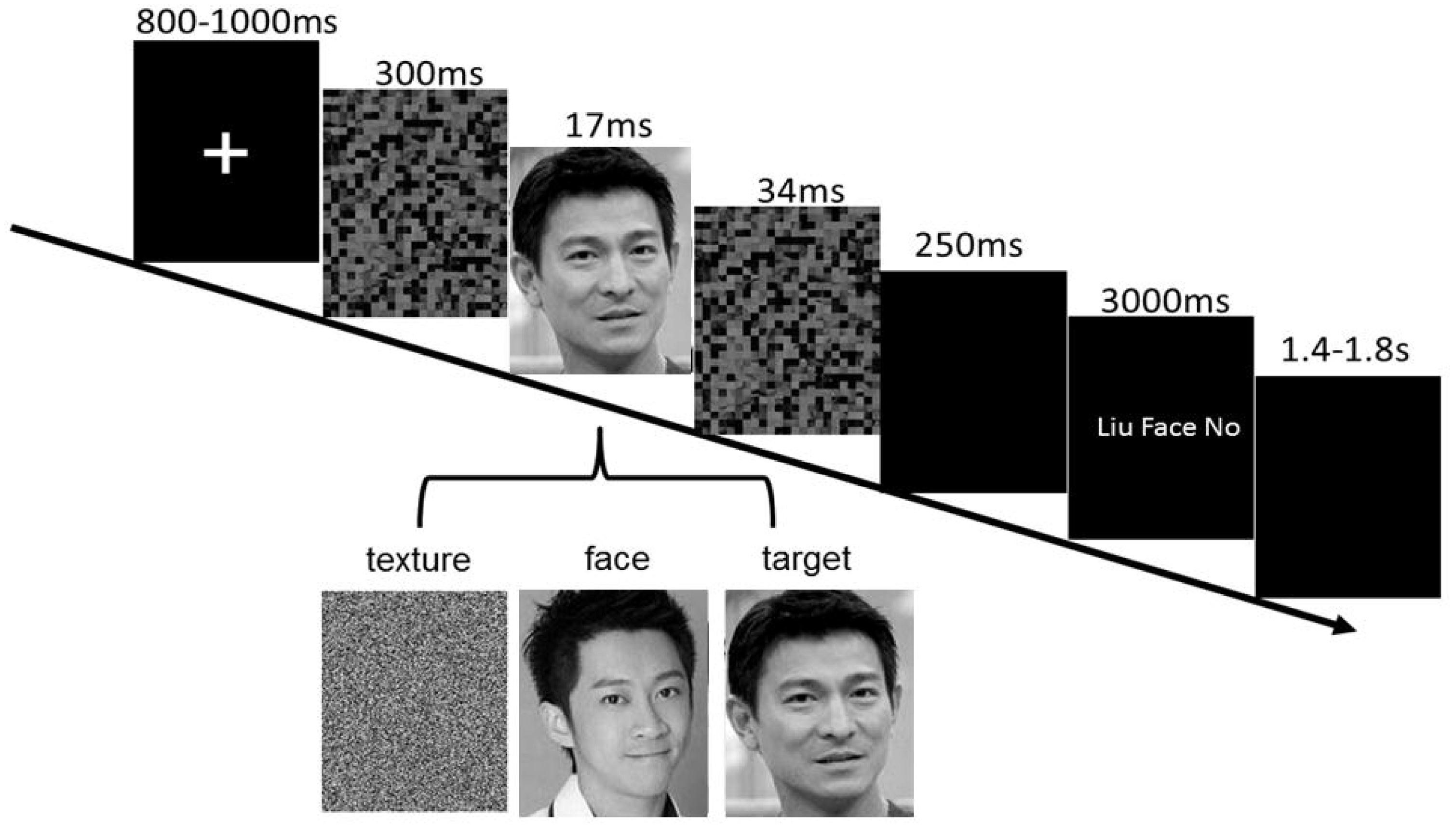
Experimental paradigm. Each trial began with the presentation of a fixation cross, followed by a forward mask, a stimulus, and a backward mask. The stimulus was either a target face, the face of another celebrity, or a texture. Participants were requested to report (1) that they could recognize the facial identity of the target stimulus, (2) that they could detect the presence of the target face but could not recognize its facial identity, or (3) that they could not see a face.

The experiment consisted of 12-14 runs. Each run consisted of 28 different target faces portraying the same celebrity, 12 other celebrity faces (30% of the face-present trials), and 17 textures (30% of the total trials). Both texture and celebrity-face trials served as catch trials and were therefore excluded from the main analysis. To ensure that participants did not randomly respond their conscious access to target identity and target configural features, we examined the percentage of the celebrity-face and texture trials in which the participants responded “Liu” (Mean = 1.06% SD(0.27)) and the percentage of the texture trials in which participants responded “Face” (Mean = 0.63% SD(0.20)). Both ratios are low and indicate that participants’ conscious access reports are not based on pure guesses. Moreover, there was no significant difference between those two ratios [paired t-test, t(12) = 1.28, p = 0.22].

To ensure a sufficient number of trials for the analysis of each condition, we adjusted mask intensities for individual participants prior to the experiment to obtain the luminance that yielded a stimulus at threshold, such that the proportion of trials in which participants consciously recognized the facial identity of the target, the proportion of trials in which participants could not recognize the identity of the target but could consciously detect the presence of a target face, and the proportion of trials in which participants could not consciously detect the presence of a target face were all above 15% (see the “Behavioral performance” section for more details). During this calibration session, the experimental paradigm was the same as described above, and each participant completed 1-3 runs.

### MEG recording and preprocessing

MEG recordings were performed using a whole-head system consisting of 157-axial-gradiometer channels (Yokogawa, Co., Tokyo). The signals were digitized at 1000 Hz and filtered with 0.3 Hz high-pass and 500 Hz low-pass cutoffs and a 60 Hz notch. To minimize head movements between runs, the participants’ head positions relative to the MEG sensors were monitored using a set of head localization coils placed at the nasion and the left and right ear canals. FieldTrip (Oostenveld et al. 2011), CircStat (Berens 2009) and MATLAB (MathWorks, Natick, MA) software were used for data preprocessing, analysis and visualization.

Continuous MEG data were segmented into 1800-ms epochs starting from 800 ms before the onset of the forward mask. Trials contaminated with eye movements, eye blinks, and muscular artifacts were rejected via visual inspection and semi-automatic functions implemented in Fieldtrip. The remaining trials were then submitted to the following analyses.

### Event-related magnetic field (ERF) analysis

The MEG data were resampled at 250 Hz and digitally filtered with a 50 Hz low-pass filter. Notably, these specific downsampling and low-pass filtering procedures were applied for the ERF analysis only. Next, the data were averaged across trials for each experimental condition and each participant. The averaged data were baseline-corrected by subtracting the mean activity during the baseline period (300-100 ms preceding the onset of the forward mask).

### Spectral power analysis

Time-frequency representations of power and phase information in the MEG signals were computed using Morlet’s wavelets (m = 7) on every sensor, frequency (8-100 Hz, step: 2 Hz) and time point (-800-1000 ms, step: 10 ms) in each trial. For spectral power, the activities were averaged across trials for each experimental condition and each participant. To compensate for 1/f decay, the averaged power activities for each data point were normalized to a baseline ranging from 350-150 ms preceding the onset of the forward mask. Normalization involved calculating the 10 log transform of the power relative to the mean baseline power on a frequency-by-frequency basis.

### Inter-trial phase coherence analysis

For phase information, the degree of phase clustering (or phase locking) in response to the target was assessed by inter-trial coherence (ITC, or phase-locking factor (TallonBaudry et al. 1996)). The ITC was computed by normalizing the lengths of complex vectors (derived from the wavelet transform) to a value of one and then computing their average across trials. The ITC index ranges between 0 and 1. An ITC close to 0 reflects low phase clustering (i.e., the distribution of phase angles across trials is uniform), whereas an ITC close to 1 reflects strong phase clustering (i.e., all trials exhibit the same phase).

### Phase bifurcation analysis

Given that two experimental conditions with comparable strength of phase clustering (i.e., similar ITC values) might be locked to different phase angles, a phase bifurcation index was calculated for every sensor-time-frequency point (s, t, f) to quantify any difference in the distribution of the phase angles between the two conditions (Busch et al. 2009). Rather than directly examining the phase distributions, this index serves as a more sensitive measure to mitigate the concern that associated power may obscure potential phase effects (Mathewson et al. 2009; VanRullen et al. 2011). To obtain this index, the respective ITC values of the conditions A and B were computed against the ITC value of the trials combined from both conditions as follows:

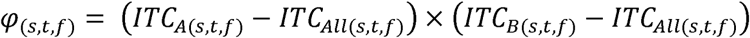

When the two conditions exhibit comparable phase clustering but different phase angles, the index is a positive value. When the degrees of phase clustering are comparable and the phases in both conditions are in a similar direction, the value is close to zero. Notably, given that a recent simulation study suggests that a variant measure, phase opposition sum (*φ* = *ITC_A_* + *ITC_B_* − 2*ITC_all_*), may provide more satisfactory outcomes (VanRullen 2016); therefore, we additionally computed the data based on this measure. The same pattern of results was still obtained.

### Bootstrap procedure

A bootstrap procedure was used to ensure that the difference in trial numbers between experimental conditions did not bias the measurement of spectral power, ITC, or the phase bifurcation index. Specifically, the trials were randomly selected without replacement from the condition with a greater number of trials, such that the trial numbers were matched between the two conditions being compared. This procedure was repeated 2000 times to generate a distribution of power, ITC or the phase bifurcation index for every sensor-time-frequency point in the condition with more trials. The obtained distribution was then characterized by its mean, which was later used in the cluster-based permutation analysis.

Notably, according to our *a priori* hypothesis, the data were computed only for the alpha band during the analysis of phase bifurcation. In addition, the experimentally obtained index was tested against the mean of a surrogate distribution under a null hypothesis (Busch et al. 2009). The reason that the index cannot be directly compared with zero is because ITC values depend on the number of trials and thereby, even for a uniform phase distribution, the expected value is not zero. The null distribution was generated by pooling all trials from the conditions, randomly assigning an equal number of trials to one of the two conditions and then recalculating the bifurcation index 2000 times.

### Cluster-based permutation test

To determine whether the data differed significantly between experimental conditions, cluster-based permutation tests were conducted (Maris and Oostenveld 2007). This statistical test does not require specific assumptions about the shape of the population distribution, and it controls for multiple comparison problems. In these tests, the experimental differences were quantified by means of paired t-tests for every sensor-time-frequency sample. The samples with t values exceeding the threshold (p < 0.05) were then clustered in connected sets based on spatial, temporal or frequency adjacency. The cluster with the maximum sum of t values was used as a test statistic. A distribution was then generated by randomly permuting the data across the conditions for each participant and then recalculating the test statistic 1000 times using a Monte Carlo estimate. Finally, p values were determined by evaluating the proportion of the distribution resulting in a test statistic larger than the observed statistic.

## RESULTS

### Behavioral performance

Because our goal was to examine how conscious access to different levels of target contents alternated over trials, the catch (i.e., non-target face and texture stimuli) trials were discarded and the target trials were sorted into the three following experimental conditions based on participants’ reports: (1) *face identification hit* (*FI)*: trials in which participants consciously recognized the facial identity of the target (mean number of trial = 55 (SD 6)); (2) *face detection hit* (*FD)*: trials in which participants could not recognize the identity of the target but could consciously detect the presence of a target face (193 (SD17)); and (3) *face miss (FM)*: trials in which participants could not consciously detect the presence of a target face (96 (SD14)). Behaviorally, the first trial type is operationally defined as reflecting conscious access to facial identity, whereas the second one reflects conscious access to facial configural features.

The participants had an average identification rate of 16.31% (SD 1.94) and a detection rate of 54.21% (SD 4.68). Here, the identification and detection rates are defined as the proportion of the FI and FD trials, respectively, to the sum of the FI, FD and FM trials. The detection rates were significantly higher than the identification rates [paired t-test, t(12) = 6.20, p < 0.001], indicating that participants were more likely to consciously access facial configural features than facial identity. To probe whether conscious access to deep-level target contents required more processing, reaction time (RT) data were examined. The analyses did not reveal any significant RT difference between the FI (Mean = 944.29 ms (SD 50.91) from the onset of the response window) and FD (Mean = 903.28 ms (SD 63.68)) trials [paired t-test, t(12) = 0.70, p = 0.50], or even when all three trial types were compared [FM: 944.29 ms (SD 50.91); one-way repeated measures ANOVA, F(2, 26) = 0.68, p = 0.4].

### Face-selective ERF activity

To validate the effectiveness of task manipulation and to identify an independent face-selective window for the follow-up analysis on the ERF difference between conscious face identification and conscious face detection, we first examined the presence of the face-selective M170 component in the data (Liu et al. 2002). Given that this component is more negative for face stimuli than for control non-face stimuli, we contrasted the ERF activities of the celebrity-face trials associated with “Face” responses (the F trials) with the texture trials associated with “No” responses (the T trials). As shown in Fig. 2A, a significant ERF difference was indeed observed between these two trial types (cluster-based permutation test, p = 0.038). Face-selective ERF responses were identified over the right central-parietal-temporal sensors, and the latencies ranged from 462 ms to 638 ms after first mask onset (162-338 ms after face onset). Visual scrutiny further revealed that those ERF activities exhibited a negative component that peaked at a latency of approximately 170 ms after face onset and exhibited a higher magnitude in the celebrity-face trials than in the texture trials. Overall, these patterns closely correspond to the previously described M170 component. It is worth mentioning that although the result was not significant, visual inspection revealed another positive component elicited approximately 100 ms after face onset, which is in agreement with another less frequently reported M100 face-sensitive component (Liu et al. 2002).

**Fig. 2.**
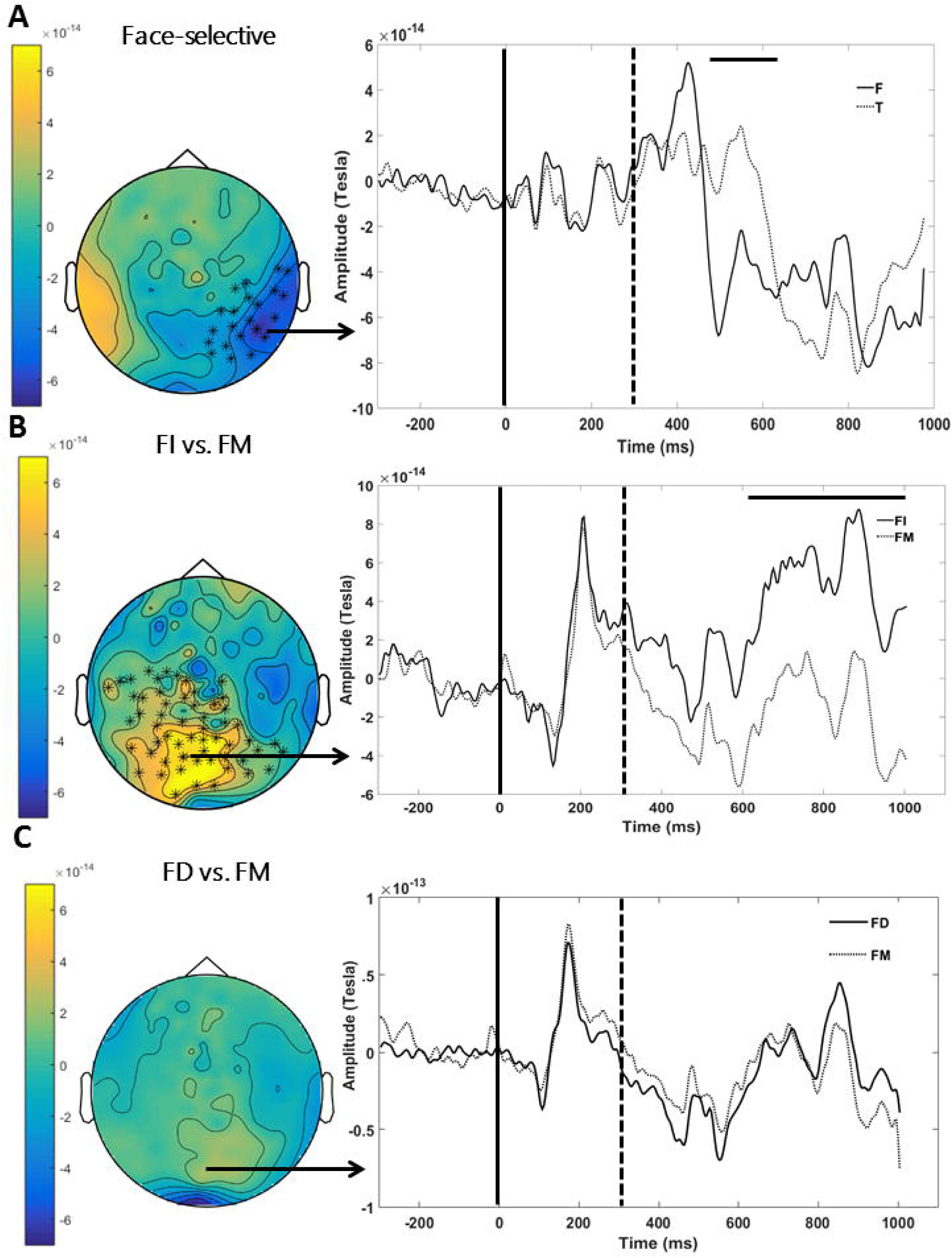
(A) The face-selective sensor-time window. Left panel: scalp topography with color scale expressing average amplitude (Tesla/cm) during the time period indicated by the horizontal bar on the right panel. The black asterisks highlight the clusters of sensors yielding significant differences. Right panel: the time course of the ERF waveform from the representative sensor AG119. The black line indicates the onset of the first mask, the dotted line indicates the onset of the stimulus, and the horizontal bar highlights the time period yielding a significant difference. (B) ERF activity during conscious face identification and (C) during conscious face detection. The conventions are the same as those in (A), except that the waveform is from sensor AG140.

### Stronger ERF activity during conscious face identification relative to conscious face detection

To investigate the neural signatures reflecting conscious access to the levels of target contents, we reasoned that differential neural responses between the FI and FM conditions would indicate conscious face identification, whereas differential responses between the FD and FM conditions would indicate conscious face detection. As in previous studies, we first examined whether the ERF responses in the previously identified face-selective window in Fig. 2A reflected those differences. For this analysis, the responses in the window were extracted and averaged for the FI, FD, and FM conditions. During conscious face identification, we observed stronger negative ERF activation in FI than in FM [paired t-test, t(12) = 2.45, p = 0.031]. In contrast, no significant ERF activity was associated with conscious face detection, as indicated by comparable responses in the FD and FM conditions [t(12) = 0.87, p = 0.40].

To provide a more rigorous measure and mitigate multiple comparison problems, cluster-based permutation tests were conducted to search for ERF activities that reflected levels of conscious access based on all sensor-time points. As seen in Fig. 2B, these analyses revealed significantly higher ERF activities in the FI than in the FM condition over the central-occipito-parietal sensors in the 606-1000 ms interval after first mask onset (306-700 ms after face onset; p = 0.018). Similar to the previously reported results, there was no significant difference between the FD and FM conditions (p > 0.29; Fig. 2C). However, when the analysis was restricted to the aforementioned sensor-frequency-time window as indicated in Fig. 2B, we did observe enhanced ERF activities in the FD trials relative to the FM trials [paired t-test, t(12) = 2.41, p = 0.033]. In sum, conscious face identification elicits more robust ERF activation than conscious face detection.

### Stronger phase clustering during conscious face identification relative to conscious face detection

In addition to ERF analysis, the spectral power and phase of the MEG data were characterized for every sensor-time-frequency point in each trial. Moreover, a bootstrap procedure was used to ensure that any difference in trial numbers between the two conditions being compared would not bias the following time-frequency analyses (see the “Bootstrap procedure*”* section for details). Although spectral power did not differ significantly during conscious face identification (cluster-based permutation test, p > 0.06) or conscious face detection (p > 0.20), a significant difference in phase clustering was observed for conscious face identification. Here, phase clustering was measured by the ITC index, which assessed to what extent phases were clustered in a similar direction over trials in each condition. As shown in Fig. 3A, phase was more densely concentrated in the FI trials than in the FM trials for the 10 Hz alpha frequency (cluster-based permutation test, p = 0.004). Moreover, this effect occurred primarily in the right frontal-parietal-temporal sites at approximately 240 ms to 580 ms after first mask onset (-60-280 ms after face onset). Although the FD and FM conditions were not significantly different (Fig. 3B; p = 0.31), when the analysis was restricted to the sensor-frequency-time window indicated in Fig. 3A, the ITC values in the FD trials were significantly higher than those in the FM trials [paired t-test, t(12) = 2.92, p = 0.013].

**Fig. 3.**
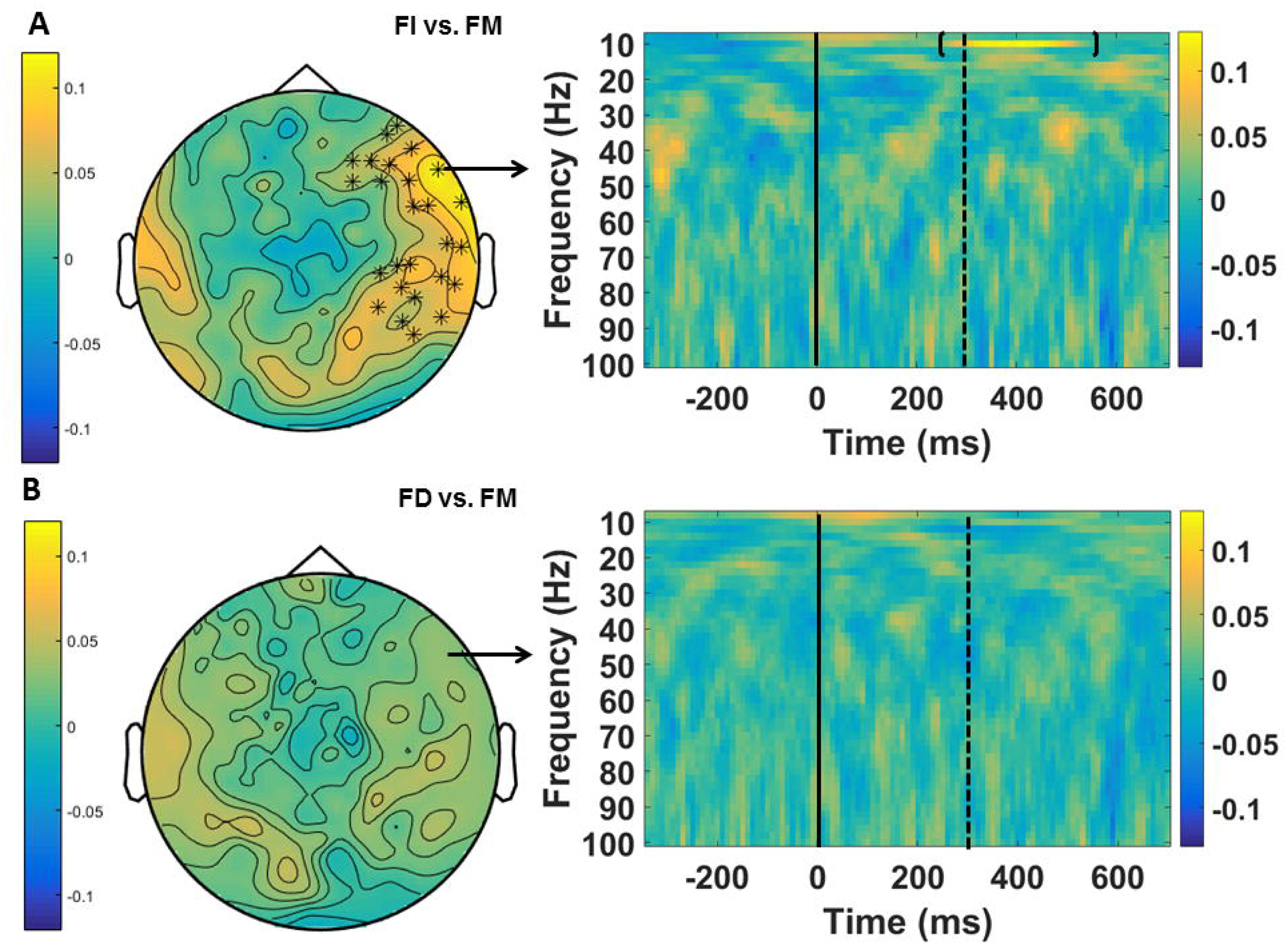
Phase clustering for (A) conscious face identification and (B) conscious face detection as revealed by the ITC measure. The color scales express the average ITC differences between (A) the FI and FM trials and (B) the FD and FM trials during the time period indicated by the horizontal bar on the right panel. The black asterisks in the scalp topographies highlight a cluster of sensors exhibiting significant differences. The right panel is a time-frequency representation of the ITC index from sensor AG98. The black line indicates the onset of the first mask, and the dotted line indicates the onset of the stimulus. The bracket highlights the time-frequency window exhibiting a significant difference.

### Distinct patterns of phase angle adjustment between conscious face identification and conscious face detection

Recently, conscious perception has been linked to the fluctuation of prestimulus phases in the alpha frequency range, with different phase angles associated with aware and unaware trials (Busch et al. 2009; Mathewson et al. 2009). Accordingly, although the previous ITC results reveal that during the prestimulus period, the two types of trials being compared (i.e., the FI and FM trial pair during conscious face identification or the FD and FM trial pair during conscious face detection) could elicit comparable phase clustering strength, it is possible that these two types of trials are locked to different phase angles. In other words, the distributions of the phase angles between the FI and FM trials or between the FD and FM trials are deviated in a similar amount from uniformity, but the direction of deviation might differ. To investigate this possibility, we did not directly assess differences in the distribution of phase angles in the alpha band (8-12 Hz) because power may obscure phase values (Mathewson et al. 2009; VanRullen et al. 2011). Instead, the phase bifurcation index in the alpha band in which power is normalized was used to examine whether those two trial types in each pair were locked to different phase directions (Busch et al. 2009; VanRullen 2016). The obtained index was then tested against shuffled data under the null hypothesis (see the “Bootstrap procedure” section for details). The positive index would represent different phase angles between trial types.

During conscious face detection, the analysis did demonstrate positive phase bifurcation indices that were significantly widespread across the brain in the 8-12 Hz frequency range and in the ‐340-80 ms time window after the onset of the first mask (Fig. 4A, cluster-based permutation test, p < 0.001). For conscious face identification, we also found that different phase angles locked to the FI and FM trials in the 8-12 Hz frequency band (Fig. 4B). However, the pattern was distinct from that observed for conscious face detection, as the phase effect occurred primarily at the anterior sensors between ‐170-110 ms following the onset of the first mask (p < 0.001). To further explore whether the obtained phase effects were indeed distinct between the two forms of conscious access, we additionally examined phase bifurcation between the FI and FD trials. As shown in Fig. 4C, the phases in these two trial types also clustered in different directions. This significant phenomenon was observed between 8-12 Hz and between ‐350-170 ms after the onset of the first mask. Overall, our observation indicates that the conscious identification and detection of the target faces could be manifested by differential patterns of phase adjustment, even prior to the onset of the first mask.

**Fig. 4.**
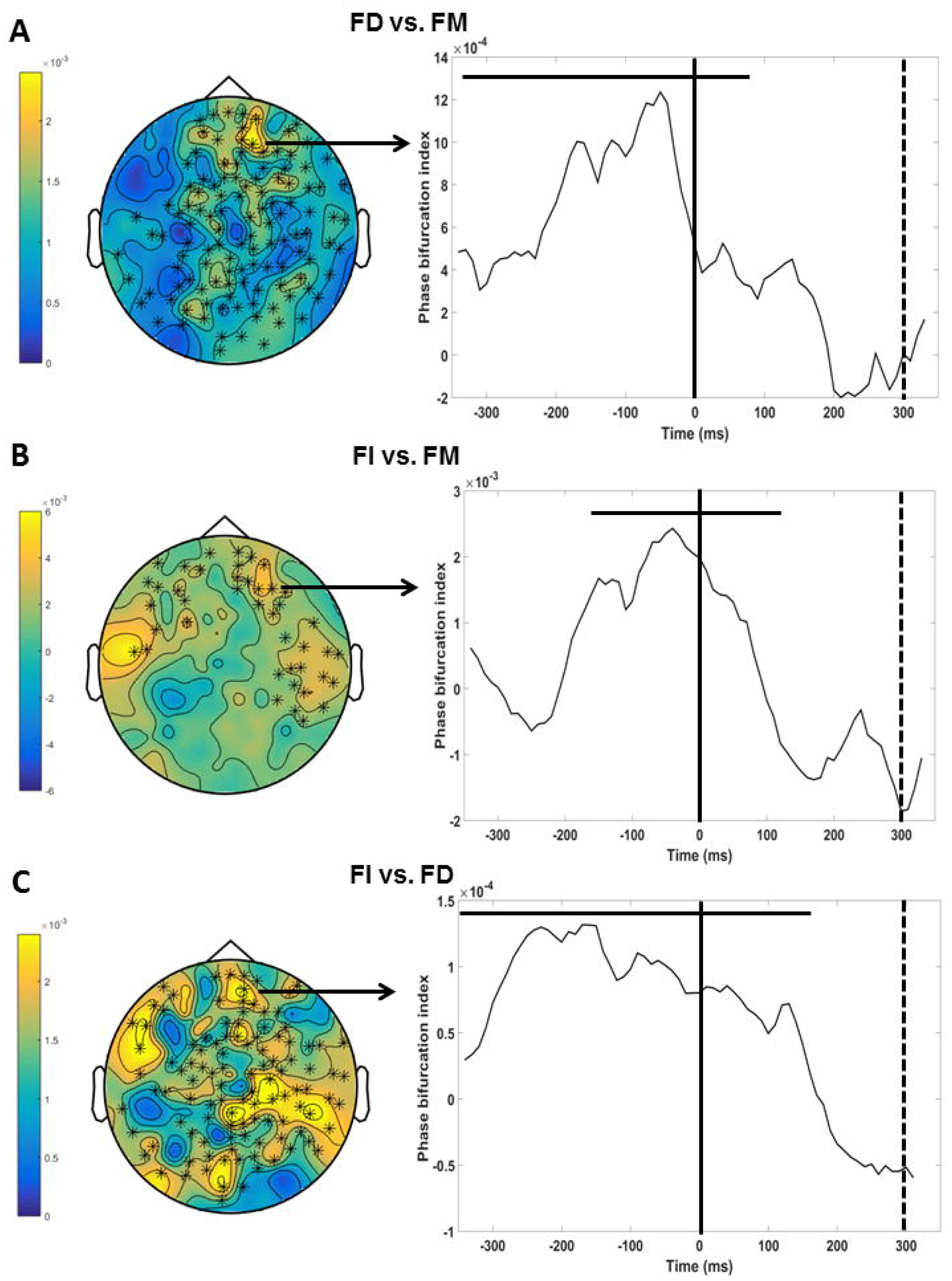
Phase bifurcation for (A) conscious face detection, (B) conscious face identification, and (C) between conscious face identification and detection. The color scales express the average phase bifurcation index derived from (A) the FD and FM trials, (B) the FI and FM trials, or (C) the FI and FD during the time period indicated by the horizontal bar on the right panel. The black asterisks in the scalp topographies highlight the clusters of sensors exhibiting significant differences. The right panels indicate the time course of the index obtained from sensors (A) AG98, (B) AG92 and (C) AG98 after collapsing across frequencies. The black lines indicate the onset of the first mask, and the dotted lines indicate the onset of the stimulus. The horizontal bars indicate the time periods exhibiting significant differences.

To substantiate the above findings, we first verified that the obtained results were not spurious, as Rayleigh’s tests were adopted to ensure that for every participant, phase distributions were not significantly deviated from uniformity when the FI and FM trials were combined (all ps > 0.25), when the FD and FM trials were combined (all ps > 0.085, except for one participant: p = 0.04) or when the FI and FD trials were combined (all ps > 0.089). In addition, we confirmed that the ITC values were indeed comparable between the given two trial types being compared in all the sensor-time-frequency windows indicated in Fig. 4 [FI vs. FM: paired t-test, t(12) = 0.19, p = 0.86; FD vs. FM: t(12) = 090, p = 0.38; FI vs. FD: t(12) = 1.53, p = 0.15]. Furthermore, current time-frequency analysis relied on convolving MEG signals with a wavelet function, which is extended in time. To ensure that the data collected after stimulus onset did not affect the prestimulus effects and vice versa, we repeated the analysis with a shorter wavelet cycle (m = 3). The main results of prestimulus phase adjustment effects and peristimulus phase clustering effects were still obtained.

## DISCUSSION

As revealed by a series of neural events, conscious access to different levels of stimulus contents (i.e., facial identity and facial configural features) shows several quantitative and qualitative distinctions. First, before the target face was actually presented, different alpha phase angle adjustment was found to link to conscious face identification and conscious face detection. The sensor-time evolution of this phenomenon also differed. Next, divergent conscious processes continued to unfold during target onset. Just before the targets were presented, the alpha phases became increasingly clustered in a similar direction across trials for conscious face identification relative to conscious face detection. After target onset, stronger ERF activity was elicited by conscious face identification as opposed to conscious face detection.

### Qualitative differences in prestimulus phase effects between conscious access to levels of facial contents

Growing evidence has addressed the importance of the ongoing phase in informing cognitive processes (Sauseng and Klimesch 2008; VanRullen et al. 2011). In particular, the prestimulus alpha phase may influence the subsequent conscious detection of near-threshold visual stimuli (Busch et al. 2009; Mathewson et al. 2009). In these studies, the aware and unaware stimuli were locked at different prestimulus phase angles. Because alpha phases may represent cortical excitability (Palva and Palva 2011), the prestimulus phases of alpha oscillations have been interpreted to reflect how the brain state, just preceding the event, oscillates between low and high excitable states in successive cycles and in turn, determines whether a stimulus is consciously perceived or not. In other words, near-threshold stimuli preceded by alpha phases indexing high excitability are more easily consciously perceived, whereas stimuli preceded by alpha phases indexing low excitability are less likely to be consciously perceived.

Consistent with this idea, the present study observed a similar role for alpha phase coding during conscious perception. We provide additional findings that prestimulus phase effects are present not only during conscious access at a basic level (i.e., facial configural features) but also at a deep level (i.e., facial identity). However, the pattern of the phase effect during conscious face identification differs from that observed during conscious face detection in a qualitative manner. First, the sensor-time evolution of the prestimulus phase effects in these two conscious processes differs. Most importantly, identified, detected and missed target faces are locked to different alpha phase angles. The latter finding suggests that conscious access to levels of facial contents could be similarly interpreted as depending on prestimulus alpha cycles, oscillating among phases of an optimal excitable state wherein the target identity could be consciously identified, phases of an intermediate excitable state wherein the targets could be consciously detected but not identifiable, and phases of a low excitable state wherein the targets could not be perceived.

### Quantitative differences in peristimulus phase clustering between conscious access to levels of facial contents

When stimuli are about to be presented, our results further reveal that phase clustering to target faces is relatively enhanced early at the alpha band to enable conscious access to facial identity rather than facial configural features. This finding concurs with a previous report indicating that alpha phase clustering to consciously perceived stimuli can be observed as early as stimulus onset (Palva et al. 2005). Because focused attention may elicit large alpha phase clustering just before or during stimulus processing (Hanslmayr et al. 2005; Yamagishi et al. 2008), our result may reflect a potential link between temporal attention and conscious access at a high level. Particularly, it is conceivable that participants were unlikely to keep a constant attentional state throughout the whole experiment. This observation might indicate their fluctuations of attention from trial to trial, such that in the FI trials, focused attention happened to be strong and in turn boosted access to high-level conscious content. In support of this idea, when deploying attention on a given location, evidence has shown that visual or conscious performance is improved by enhancing the perceptual representation of incoming stimuli (Hsu et al. 2011; Prinzmetal et al. 2005), possibly via a change in the visual system’s effective spatial resolution (Anton-Erxleben and Carrasco 2013).

### Quantitative differences in poststimulus ERF activities between conscious access to levels of facial contents

The current results concurs with prior research (Tanskanen et al. 2007), showing the involvement of the M170 component in conscious face identification. Despite some previous findings (Navajas et al. 2013; Sandberg et al. 2013), this observation indicates that conscious access to facial identify may occur at an early stage (See more discussion in the following section). We also identified a stronger late positive ERF activation during conscious face identification (Genetti et al. 2009) and conscious face detection (Rodriguez et al. 2012), with stronger magnitude in the former. The sensor-time pattern of this component closely corresponds to so-called late positivity (LP) in perceptual awareness, which predominantly localizes at the parietal and central sites and begins in the P300 time window at approximately 300-400 ms after stimulus presentation (Koivisto and Revonsuo 2010; Railo et al. 2011). However, it remains unclear whether LP is a direct correlate of conscious perception (Liddell et al. 2004; Sergent et al. 2005) or the consequences of post-awareness cognitive processes, such as working-memory maintenance (Aru et al. 2012). Despite this controversy, the LP observed in this study may suggest that a distinct stage of late processing is involved in mediating conscious access to facial identity.

### Distinct but associated neural mechanisms underlying conscious access to levels of stimulus contents

The present study investigates how conscious access to the existing levels of stimulus contents embedded in the target stimuli alternates over trials within a single experimental design. We demonstrate that quantitatively and qualitatively distinct stages of conscious processing were recruited for accessing facial identity and facial configural features. On the one hand, the neural activities underlying conscious access to levels of stimulus contents diverge even before stimulus onset. These findings suggest that different levels of stimulus contents may not be simultaneously accessed and then resolved at a later processing stage. Rather, a specific level of upcoming conscious content is separately tracked and partly determined early before stimulus is presented. Although counterintuitive, a similar account has previously proposed that stimulus contents at each level are independently consciously accessed, in the sense that conscious access to deep-level stimulus contents can occur without accessing basic-level contents and vice versa (Kouider et al. 2010).

On the other hand, our results reveal early phase clustering and late LP activity during both conscious face identification and conscious face detection, despite weak magnitude in the latter. This overlapping pattern of neural signatures suggests that conscious access to levels of stimulus contents may rely on an associated mechanism in which conscious access to deep-level contents is built upon conscious access to basic-level contents. Accordingly, to consciously access deep-level stimulus contents, early enhanced focused attention, as marked by phase clustering, and late cognitive reflection, as marked by LP activity, might be required for further processing.

In contrast to the previous studies which indicate that conscious face detection precedes conscious face identification, the present results show an early neural signature (i.e., ITC and M170 enhancement) responsible for conscious face identification after stimulus onset. This discrepancy may reflect different analysis approaches. For example, even-related analysis is primarily adopted in previous studies, whereas our findings concerning phase information are based on time-frequency analysis and a different statistical analysis is employed to mitigate multiple comparison problems, which might also explain why relatively weak neural responses to conscious face detection cannot be properly characterized. However, the discrepancy may also reflect different task requirements. In the current design, conscious face identification and detection were investigated within a single paradigm and conscious access at different levels was framed based on participants’ response choices. Accordingly, is it possible that these choices did not genuinely reflect levels of conscious access, but processing bias due to the nature of the task? For example, participants were aware which target celebrity that they should expect. Because of such bias (Melloni et al. 2011), conscious face identification elicited stronger ITC and M170 responses than conscious face detection and the responses occurred early right after stimulus onset. Although this possibility cannot be completely ruled out, we suggest that the overall findings are not fully compatible with this alternative explanation. First, it is worth emphasizing that catch trials were included in the design so that participants did not have any prior knowledge regarding which type of stimulus was about to be presented. In addition, participants were instructed to report their conscious experience of the stimuli rather than searching for target identity in the task. More importantly, our analysis does not reveal any RT differences among response choices. In addition, participants produced few false identifications and detections during catch trials, and these two types of false responses did not significantly differ (see the “Procedure” section). These analyses collectively reveal that general processing bias may not exist when selecting the choice “Liu” and “Face”. Notably, expectation lowers the threshold of conscious perception (Melloni et al. 2011) and thereby high identification rates should be obtained if expectation does exist and benefits conscious face identification. However, our behavioral data show that when facing an incoming stimulus, participants were better at consciously accessing facial configural features than facial identity, as indicated by higher detection rates. Taken together, we suggest that “task advantage” specific to conscious face identification is unlike to be present in this study.

Nevertheless, in the current setting, only one target celebrity was used throughout the experiment. Although this strategy aims to avoid introducing additional noise, given that different target identities require differential mask intensities to obtain the luminance that yields stimuli at threshold, our approach inevitably limits the generalization of our findings to a broader domain of stimulus processing. Additionally, a more sensitive response scale, such as refined and broad levels of response choices, should be incorporated in future investigation to better understand the neural mechanism underlying levels of conscious face perception.

### Conclusion

Beyond the controversy surrounding whether early or late neural responses are responsible for conscious face detection or identification, we suggest that prestimulus activities together with stimulus-related activities may operate in combination to determine which level of stimulus contents can be consciously accessed. This idea is in line with recent accumulating evidence showing that perceptual processing depends not only on stimuli themselves but also on the state of rhythmic brain activity that precedes the upcoming stimuli (Britz and Michel 2011 for a review). Thus, conscious access may be better construed as comprising various forms that depend on the level of stimulus contents accessed. We believe that this view will provide a more solid basis for understanding the nature of conscious access in future studies.

## ACKNOWLEDGMENT

This work was supported by the Ministry of Science of Taiwan, R.O.C. (NSC 102-2410-H-004-045-MY2 and NSC 104-2410-H-004-049). The authors thank Catherine Tallon-Baudry and Andy Young for their comments on an early version of the manuscript.

## REFERENCES

Anton-Erxleben K, and Carrasco M. Attentional enhancement of spatial resolution: linking behavioural and neurophysiological evidence. Nat Rev Neurosci 14: 188–200, 2013.

Aru J, Bachmann T, Singer W, and Melloni L. Distilling the neural correlates of consciousness. Neurosci Biobehav Rev 36: 737–746, 2012.

Baars BJ. The conscious access hypothesis: origins and recent evidence. Trends Cogn Sci 6: 47–52, 2002.

Bentin S, Allison T, Puce A, Perez E, and McCarthy G. Electrophysiological studies of face perception in humans. J Cogn Neurosci 8: 551–565, 1996.

Berens P. CircStat: a MATLAB toolbox for circular statistics. J Stat Softw 31: 1–21, 2009.

Britz J, and Michel CM. State-dependent visual processing. Front Psychol 2: 370, 2011.

Bruce V, and Young A. Understanding face recognition. Brit J Psychol 77: 305–327, 1986.

Busch NA, Dubois J, and VanRullen R. The phase of ongoing EEG oscillations predicts visual perception. J Neurosci 29: 7869–7876, 2009.

Campana F, and Tallon-Baudry C. Anchoring visual subjective experience in a neural model: The coarse vividness hypothesis. Neuropsychologia 51: 1050–1060, 2013.

Dehaene S, and Changeux JP. Experimental and theoretical approaches to conscious processing. Neuron 70: 200–227, 2011.

Genetti M, Khateb A, Heinzer S, Michel CM, and Pegna AJ. Temporal dynamics of awareness for facial identity revealed with ERP. Brain Cognition 69: 296–305, 2009.

Hanslmayr S, Klimesch W, Sauseng P, Gruber W, Doppelmayr M, Freunberger R, and Pecherstorfer T. Visual discrimination performance is related to decreased alpha amplitude but increased phase locking. Neurosci Lett 375: 64–68, 2005.

Haxby JV, Hoffman EA, and Gobbini MI. The distributed human neural system for face perception. Trends Cogn Sci 4: 223–233, 2000.

Hochstein S, and Ahissar M. View from the top: hierarchies and reverse hierarchies in the visual system. Neuron 36: 791–804, 2002.

Hsu SM, George N, Wyart V, and Tallon-Baudry C. Voluntary and involuntary spatial attentions interact differently with awareness. Neuropsychologia 49: 2465–2474, 2011.

Koivisto M, Grassini S, Salminen-Vaparanta N, and Revonsuo A. Different electrophysiological correlates of visual awareness for detection and identification. J Cognitive Neurosci 29: 1621–1631, 2017.

Koivisto M, and Revonsuo A. Event-related brain potential correlates of visual awareness. Neurosci Biobehav Rev 34: 922–934, 2010.

Kouider S, de Gardelle V, Sackur J, and Dupoux E. How rich is consciousness? The partial awareness hypothesis. Trends Cogn Sci 14: 301–307, 2010.

Kouider S, and Dehaene S. Levels of processing during non-conscious perception: a critical review of visual masking. Philos T R Soc B 362: 857–875, 2007.

Liddell BJ, Williams LM, Rathjen J, Shevrin H, and Gordon E. A temporal dissociation of subliminal versus supraliminal fear perception: an event-related potential study. J Cogn Neurosci 16: 479–486, 2004.

Liu J, Harris A, and Kanwisher N. Stages of processing in face perception: an MEG study. Nat Neurosci 5: 910–916, 2002.

Maris E, and Oostenveld R. Nonparametric statistical testing of EEG‐ and MEG-data. J Neurosci Meth 164: 177–190, 2007.

Mathewson KE, Gratton G, Fabiani M, Beck DM, and Ro T. To see or not to see: prestimulus alpha phase predicts visual awareness. J Neurosci 29: 2725–2732, 2009.

Maurer D, Le Grand R, and Mondloch CJ. The many faces of configural processing. Trends Cogn Sci 6: 255–260, 2002.

Melloni L, Schwiedrzik CM, Muller N, Rodriguez E, and Singer W. Expectations change the signatures and timing of electrophysiological correlates of perceptual awareness. J Neurosci 31: 1386–1396, 2011.

Navajas J, Ahmadi M, and Quiroga RQ. Uncovering the mechanisms of conscious face perception: A single-trial study of the N170 responses. J Neurosci 33: 1337–1343, 2013.

Oostenveld R, Fries P, Maris E, and Schoffelen JM. FieldTrip: open source software for advanced analysis of MEG, EEG, and invasive electrophysiological data. Comput Intel Neurosc 2011: 156869, 2011.

Palva S, Linkenkaer-Hansen K, Naatanen R, and Palva JM. Early neural correlates of conscious somatosensory perception. J Neurosci 25: 5248–5258, 2005.

Palva S, and Palva JM. Functional roles of alpha-band phase synchronization in local and large-scale cortical networks. Front Psychol 2: 204, 2011.

Prinzmetal W, McCool C, and Park S. Attention: reaction time and accuracy reveal different mechanisms. J Exp Psychol Gen 134: 73–92, 2005.

Railo H, Koivisto M, and Revonsuo A. Tracking the processes behind conscious perception: a review of event-related potential correlates of visual consciousness. Conscious Cogn 20: 972–983, 2011.

Rodriguez V, Thompson R, Stokes M, Brett M, Alvarez I, Valdes-Sosa M, and Duncan J. Absence of face-specific cortical activity in the complete absence of awareness: Converging evidence from functional magnetic resonance imaging and event-related potentials. J Cognitive Neurosci 24: 396–415, 2012.

Sandberg K, Bahrami B, Kanai R, Barnes GR, Overgaard M, and Rees G. Early visual responses predict conscious face perception within and between subjects during binocular rivalry. J Cognitive Neurosci 25: 969–985, 2013.

Sauseng P, and Klimesch W. What does phase information of oscillatory brain activity tell us about cognitive processes? Neurosci Biobehav Rev 32: 1001–1013, 2008.

Sergent C, Baillet S, and Dehaene S. Timing of the brain events underlying access to consciousness during the attentional blink. Nat Neurosci 8: 1391–1400, 2005.

Tallon-Baudry C, Bertrand O, Delpuech C, and Pernier J. Stimulus specificity of phase-locked and non-phase-locked 40 Hz visual responses in human. J Neurosci 16: 4240–4249, 1996.

Tanskanen T, Nasanen R, Ojanpaa H, and Hari R. Face recognition and cortical responses: Effect of stimulus duration. Neuroimage 35: 1636–1644, 2007.

VanRullen R. How to evaluate phase differences between trial groups in ongoing electrphysiological signals. Front Neurosci-Switz 10: 426, 2016.

VanRullen R, Busch NA, Drewes J, and Dubois J. Ongoing EEG phase as a trial-by-trial predictor of perceptual and attentional variability. Front Psychol 2: 60, 2011.

Willenbockel V, Sadr J, Fiset D, Horne GO, Gosselin F, and Tanaka JW. Controlling low-level image properties: the SHINE toolbox. Behav Res Methods 42: 671–684, 2010.

Windey B, Gevers W, and Cleeremans A. Subjective visibility depends on level of processing. Cognition 129: 404–409, 2013.

Yamagishi N, Callan DE, Anderson SJ, and Kawato M. Attentional changes in pre-stimulus oscillatory activity within early visual cortex are predictive of human visual performance. Brain Res 1197: 115–122, 2008.

